# SPsimSeq: semi-parametric simulation of bulk and single cell RNA sequencing data

**DOI:** 10.1101/677740

**Authors:** Alemu Takele Assefa, Jo Vandesompele, Olivier Thas

## Abstract

SPsimSeq is a semi-parametric simulation method for bulk and single cell RNA sequencing data. It simulates data from a good estimate of the actual distribution of a given real RNA-seq dataset. In contrast to existing approaches that assume a particular data distribution, our method constructs an empirical distribution of gene expression data from a given source RNA-seq experiment to faithfully capture the data characteristics of real data. Importantly, our method can be used to simulate a wide range of scenarios, such as single or multiple biological groups, systematic variations (e.g. confounding batch effects), and different sample sizes. It can also be used to simulate different gene expression units resulting from different library preparation protocols, such as read counts or UMI counts.

**Availability and implementation:** The R package and associated documentation is available from https://github.com/CenterForStatistics-UGent/SPsimSeq.

**Supplementary information:** Supplementary data are available at *bioR*χ*iv* online.

## Introduction

The number of computational tools for the analysis of bulk and single cell RNA sequencing (scRNA-seq) data is growing rapidly [8]. Several methods have been introduced for a single task, e.g. testing for differential gene expression (DGE). These tools typically pass through an evaluation process, often focusing on false discovery rate control and sensitivity. While such an evaluation often relies on simulated data with a built-in truth, to realistically assess the performance of these data analysis tools, the simulated data must faithfully recapitulate the data characteristics of real data [6, 5].

Various methods have been proposed for simulating either bulk or single cell RNA-seq data. The starting point is typically a distributional assumption of the gene expression data, for example the (zero inflated) negative binomial distribution [7]. While these parametric simulation methods are flexible and allow simulating various scenarios by generating synthetic data with good fit to the real data [5], such strong distributional assumptions do not hold in general. Due to the intrinsic biological variability and technical noise, scRNA-seq data sometimes show multimodal distributions [2]. There are also fully non-parametric approaches that employ subsampling from real data [3]. Although non-parametric simulators generate realistic synthetic data, they have limited flexibility and require a large source dataset to subsample from [1].

Here, we present a new simulation procedure for simulating bulk and single cell RNA-seq data. It is designed to maximally retain the characteristics of real RNA sequencing data with reasonable flexibility to simulate a wide range of scenarios. In a first step, the logarithmic counts per million (log-CPM) values from a given real data set are used for semi-parametrically estimating gene-wise distributions. This method is based on a fast log-linear model estimation approach developed by [4]. Arbitrarily large datasets, with realistically varying library sizes, can be sampled from these distributions. Our method has an additional step to explicitly account for the high abundance of zero counts, typical for scRNA-seq data. This step models the probability of zero counts as a function of the mean expression of the gene and the library size (read depth) of the cell (both in log scale). Zero counts are then added to the simulated data such that the observed relationship (zero probability to mean expression and library size) is maintained. In addition, our method simulates DGE by separately estimating the distributions of the gene expression from the different populations (for example treatment groups) in the source data, and subsequently sampling a new dataset from each group.

Our simulation procedure enables benchmarking of statistical and bio-informatics tools with realistic simulated data. In the result section, we demonstrate that the simulated data from our method retains the characteristics of the source data in terms of variability, distribution of mean expression, fraction of zero counts, and the relationship to each other (Figure 1). The details of the procedures and implementations can be found in the supplementary file. Data simulated with our procedure are compared with the original real source data and with data simulated with the parametric Splat procedure [7], which uses a gamma-Poisson hierarchical model (*splatter* R Bioconductor package, version 1.6.1, [7]).

**Figure 1:**
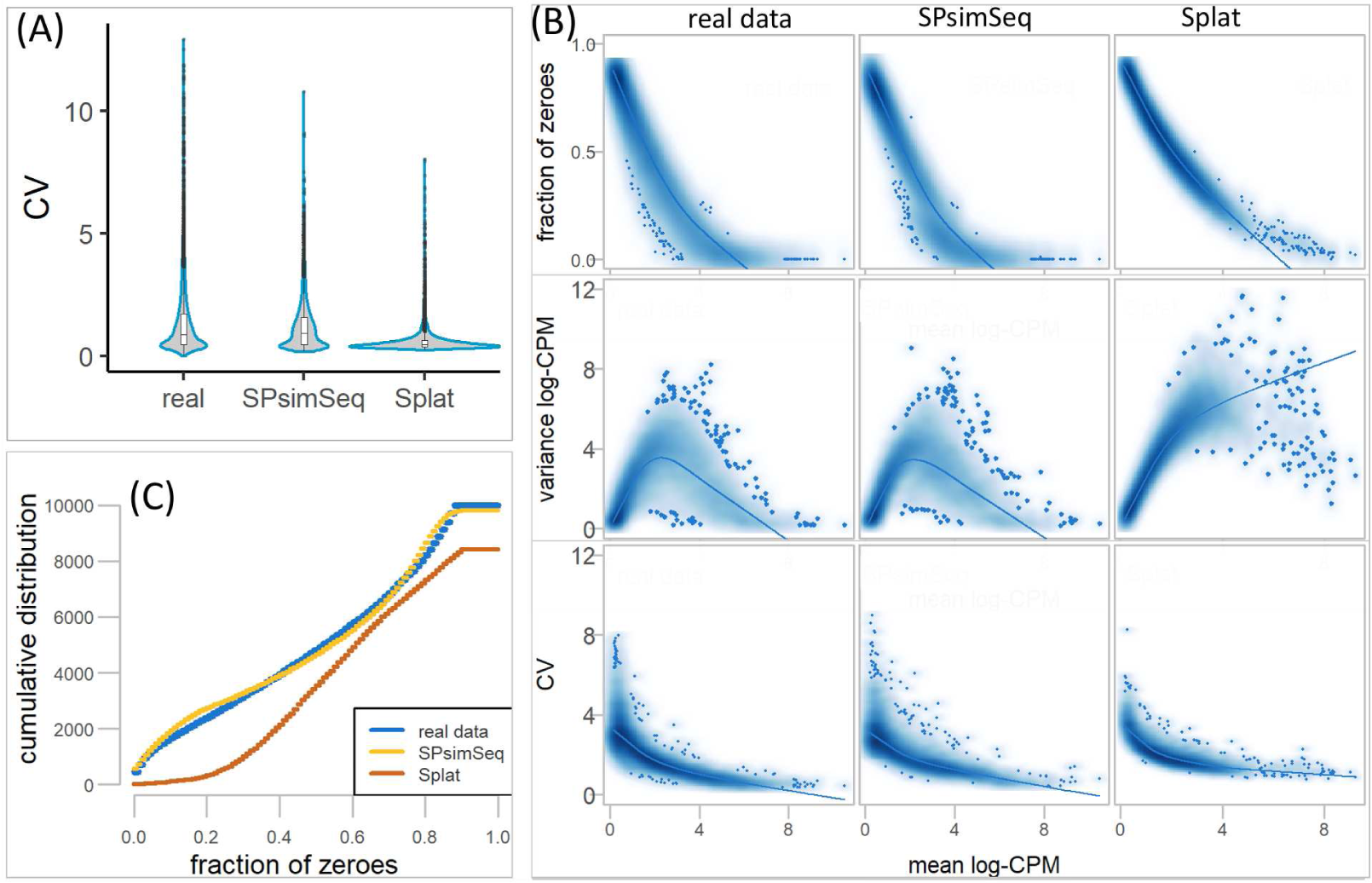
(A) Distribution of the coefficients of variations (CV) from the real and simulated (SPsimSeq and Splat) bulk RNA-seq datasets. (B) The relationship between the gene specific mean expression (in log-CPM) and three characteristics (fraction of zeroes, variance, and CV of each gene) from the real and simulated scRNA-seq datasets (read-counts). The curves show the smoothed relationship using *LOESS* regression. (C) The cumulative distribution of fraction of zero counts per gene.

## Results

Using three different source RNA-seq datasets (one bulk and two single cell) we benchmarked the novel SPsimSeq simulation method. In particular, we compared the simulated data (using SPsimSeq and Splat) with the real data with respect to various gene and sample (cell) level characteristics as used by [5] and [7]. To simulate bulk RNA-seq data using Splat, we disabled its feature for adding dropouts (*dropout.type=”none”*), which is specifically designed for scRNA-seq data simulation. The results generally show that our simulation procedure sufficiently captured the properties of the real data both for bulk and single cell RNA-seq (Figure 1 and supplementary file). The coefficients of variation, variability, distribution of mean expression and fraction of zero counts (per gene and sample/cells) in SPsimSeq simulated data resemble that of the real datasets. Compared with Splat, SPsim-Seq generates more realistic data with respect to the majority of the considered metrics. In the supplementary file, we present the detailed benchmarking results including the application of SPsimSeq for simulating scRNA-seq data with read-counts and UMI-counts (unique molecular identifier).

