## supplementary file for "SPsimSeq: semi-parametric simulation of bulk and single cell RNA sequencing data"

*February 21, 2019*

**References** **17**

#### 1 Semi-parametric simulation

The SPsimSeq simulation procedure involves two major steps: (1) density estimation for the observed read counts in a source RNA-seq dataset (bulk or single cell), and (2) simulate new count data from the estimated densities. It also involves additional steps, depending on the required simulation design. For example, simulation of single cell RNA-seq (scRNA-seq) involves an additional step to simulate the inflation of zeroes, which is an essential characteristic of scRNA-seq data. Simulation of data from multiple batches (technical batch or subject batch), involves an extra step to retain the effect of the batch in the simulated data. Simulation of differential expression (DE) also requires extra steps prior to density estimation. In the subsequent sections, we describe these steps in detail.

Let us denote the observed read counts in the source data by a $G \times n$ matrix $\mathbf{Y}$, where $G$ is the number of genes and $n$ is the total number of samples (in bulk setting) or cells (in single cell setting). Let $Y_{gij}$ be the element of the matrix $\mathbf{Y}$ for gene $g = 1, \ldots, G$, from cell $i = 1, \ldots, n_j$, of batch $j = 1, \ldots, r$, such that $n = \sum_{j=1}^{r} n_j$. For scRNA-seq data, batch can be the sample (e.g. subject or tissue) from which cells are extracted if multiple samples are used.

Density estimation and data simulation are described for a collection of cells/samples from a single batch, and later we combine them for every batch in such a way that the underlying batch effect is conserved (if required). Therefore, we drop the index $j$ and use the notation $Y_{gi}$ for the read counts of gene $g = 1, \ldots, G$, in sample/cell $i = 1, \ldots, n$.

The simulation method starts from the log-counts per millions of reads (CPM),

$$C_{gi} = \log \left\{ \frac{Y_{gi}}{L_i} \times 1e^6 + v \right\}, \tag{1}$$

where $L_i$ is the library size in sample/cell $i$. Note that $C_{gi} \in \mathbb{R}$, which can be interpreted as the log relative abundance of gene $g$ in cell $i$. It is multiplied by a constant $10^6$ to interpret the values as counts per 1 millions of reads, so that all samples have exactly a library size of 1 million. A constant $v, v > 0$ is added to avoid the logarithm of 0.

##### 1.1 Construction of the probability distribution function (density)

The first step consists of estimating the density of $C_{gi}$ from the real source data using a specially designed exponential family for density estimation as described in Efron, Tibshirani, and others (1996). Construction of the density of $C_{gi}$ begins with partitioning its sample space $\mathbb{R}$ into $K$ disjoint classes. Let the $k^{th}$ class (interval) be denoted by $s_{gk}$, such that $\cup_{k=1}^{K} s_{gk} = \mathbb{R}$. Afterwards, we define the reduced data as

$$N_{gk} = \#\{C_{gi} \in s_{gk}\}. \tag{2}$$

In other words, $N_{kg}$ is the number of $C_{gi}$ that belong to class $s_{gk}$. The number of classes $K \leq n$ is choosen automatically using the Sturges' rule (Sturges 1926) implemented in *hist()* function of the *graphics* R CRAN package (R Core Team 2018). It is also possible to specify the number of classes in terms of the fraction of the sample size in the source datset (*SPsimSeq(..., w)*, for $0 \leq w \leq 1$).

It is natural to assume that $N_{kg} \sim \text{Pois}(\mu_{kg}(\boldsymbol{\beta}_g))$. The expected number of $C_{gi}$ in class $s_{gk}$, $\text{E}\{N_{kg}\} = \mu_{kg}(\boldsymbol{\beta}_g)$, can be factorised into two factors, i.e.

$$\mu_{gk}(\boldsymbol{\beta}_g) = \mu^0_{gk} \times \mu^1_{gk}(\boldsymbol{\beta}_g), \tag{3}$$

where $\mu^1_{gk}(\boldsymbol{\beta}_g) = \exp(\beta_{g0} + \beta_{g1}t_{gk} + \beta_{g2}t^2_{gk} + \dots)$ with $t_{gk}$ any point in $s_{gk}$ (often the mid point). The factor $\mu^0_{gk}$ is called the *carrier density*, which can be the kernel smoothed density estimate of $C_{gi}$ with an appropriate band width, as in Efron, Tibshirani, and others (1996). However, CPM is a non-negative continuous approximation of gene expression, which is often characterized by a log-normal distribution to allow skewness (long right tail) (Aartsen et al. 2015). Therefore, we use a normal carrier density as a starting guess for the distribution of $C_{gi}$. Thus, for a given class $s_{gk}$,

$$\mu^0_{gk} = n\text{P}(C_{gi} \in s_{gk}|\mu_{gi}, \sigma^2_{gi}) \tag{4}$$

where $n = \sum_{k=1}^{K} N_{kg}$, and $\mu_{gi}$ and $\sigma^2_{gi}$ are the mean and variance of $C_{gi}$, respectively. The factor $\mu^1_{kg}(\boldsymbol{\beta}_g)$ indicates how $\mu_{kg}(\boldsymbol{\beta}_g)$ deviates from the carrier density. This is because, if all the $\boldsymbol{\beta}_g$ parameters are zero, then $\mu_{kg}(\boldsymbol{\beta}_g)$ reduces to the carrier density. Thus, a misspecification of the carrier density will be compensated by $\mu^1_{kg}(\boldsymbol{\beta}_g)$. For example, multimodal or skewed distributions (this is often the case for single cell RNA-seq data) will be tracked by the second factor.

Next, we fit a log-linear model (GLM with Poisson family and log-link) for the expected count in class $s_{gk}$ as a function of $t_{gk}$, i.e.

$$
\begin{aligned}
\log\{\mathrm{E}\{N_{gk}\}\} = \log\{\mu_{kg}(\boldsymbol{\beta}_g)\} &= \log\{\mu^0_{gk}\} + \log\{\mu^1_{gk}(\boldsymbol{\beta}_g)\} \\
&= \log\{\mu^0_{gk}\} + \sum_{r=0}^{R} \beta_{gr} t^r_{gk},
\end{aligned} \tag{5}
$$

where, $t_{gk}$ is the mid point of the class $s_{gk}$. We fit this model with $\mu^0_{kg}$ as an offset. Once the parameters $\boldsymbol{\theta}^T_g = (\boldsymbol{\beta}^T_g,\ \mu_{C_{gi}},\ \sigma^2_{C_{gi}})$ are estimated, the estimated density becomes ($R = 4$)

$$
\hat{f}(s_{gk}; \hat{\boldsymbol{\theta}}_g) = \hat{\mu}_{kg}(\hat{\boldsymbol{\beta}}_g) = \hat{\mu}^0_{gk} \exp\left(\sum_{r=0}^{4} \hat{\beta}_{gr} t^r_{gk}\right), \tag{6}
$$

Note that a fourth degree polynomial is chosen to capture up to the fourth degree moment of the distribution (mean, variance, skewness and kurtosis). However, for some genes with a limited number of observed data, the model in (6) may not be affordable. Thus, we successively reduce the degree of the polynomial until an estimable model is found with a minimum of a first degree polynomial.

#### 1.2 Sampling from $\hat{f}(s_{gk}; \boldsymbol{\theta}_g)$

From the estimated density in (6), we sample a new set of log CPM ($C^*_{g'i'}, i' = 1, 2, \ldots, n'$ $g' = 1, 2, \ldots, G'$), where $n'$ and $G'$ are the number of cells/samples and genes in the simulated data, respectively. In particular, we make use of the probability integral transformation to generate a new set of log CPM from the distribution function $\hat{F}(s_{gk}; \boldsymbol{\theta}_g)$ of (6). That is,

$$
\hat{F}(s_{gk}; \hat{\boldsymbol{\theta}}_g) = \int_{-\infty}^{t_{gk}} \hat{f}(h; \hat{\boldsymbol{\theta}}_g) dh. \tag{7}
$$

If $S \sim \hat{F}$, then $\hat{F}(S) \sim$ Uniform$(0, 1)$. Therefore, to sample a new data set, first we draw $n'$ observations (let us denote it by $U_i, i = 1, \ldots, n'$) from a $[0, 1]$ uniform distribution. Subsequently, $C^*_{g'i'} = \mathrm{ArgMin}_{t_{gk}} \left\{ |U_i - \hat{F}(t_{gk}; \hat{\boldsymbol{\theta}}_g)| \right\}$.

The count data ($Y^*_{i'g'}$) can be obtained as

$$Y^*_{i'g'} = \frac{(e^{C^*_{i'g'}} - 1)}{1e^6} L^*_{i'}, \tag{8}$$

where $L^*_{i'}$ is the library size in cell/sample $i'$ of the simulated data, which is sampled from the distribution of library sizes ($L_i$) in the source data. However, to allow sampling with $n' > n$, we fit a log-normal distribution to the observed $L_i$, and sample $n'$ new library sizes $L^*_{i'}$ from the fitted log-normal distribution.

##### 1.3 Simulation of zero inflation for single cell RNA-seq data

Since zeroes play an important role in scRNA-Seq, we can further improve the method by modelling the distribution of the zeroes separately (as in a hurdle model). This can be accomplished by restricting the previous steps of the model to the non-zero counts ($Y_{gi} > 0 \Rightarrow C_{gi} > 0$). We model the probability of zero counts $Z_{gi} = I(Y_{gi} = 0)$ of gene $g$ in cell $i$ as a function of the log library size $L_i$ and average log CPM ($\bar{C}_g = n^{-1} \sum_{i=1}^{n} C_{gi}$) using logistic regression. That is,

$$\text{logit}\{P(Z_{gi} = 1)\} = \lambda_0 + \lambda_1 \bar{C}_g + \lambda_2 \log L_i + \lambda_3 \bar{C}_g \times \log L_i. \tag{9}$$

Next, for each simulated gene $g'$ in the simulated cell $i'$, we compute the probability of zero expression $\hat{P}(Y^*_{g'i'} = 0)$ given the mean log CPM ($\bar{C}_g$) and library size ($L^*_{i'}$). Let this probability be denoted by $p_{g'i'}$. Next, we generate $Z^*_{g'i'}$ from a Bernoulli distribution with probability $p_{g'i'}$, i.e. $Z^*_{g'i'} \sim \text{Bernoulli}(p_{g'i'})$. Finally, the simulated expression becomes

$$Y^+_{g'i'} = Y^*_{g'i'}(1 - Z^*_{g'i'}) = \begin{cases} 0 & \text{if } Z^*_{g'i'} = 1 \\ Y^*_{g'i'} & \text{if } Z^*_{g'i'} = 0. \end{cases} \tag{10}$$

##### 1.4 Simulating differential expression (DE)

With parametric simulation methods, DE is simulated by applying a fold-change factor (Zappia, Phipson, and Oshlack 2017; Vieth et al. 2017). We simulate DE by separately estimating the distributions of the gene expression from the different populations (for example treatment groups) in the source data, and subsequently sampling a new dataset from each group. We apply this step for a set of selected genes with observed fold-change at least equal to a particular threshold.

To add a fraction of DE genes, let us denote this fraction by $p_{DE}, 0 \leq p_{DE} \leq 1$ (*pDE* in the *SPsimSeq(pDE)* function) between two or more conditions (for example treatment and control),

- Step 1: It calculates the log-fold-change for each gene in the source data
- Step 2: It selects genes with estimated log-fold-change greater than a pre-defined threshold, say $F$, (see *chooseCandGenes(lfc.thrld=F)*),
- Step 3: For the selected genes in Step 2, it fits the log-linear model in (5) by adding the grouping factor as additional covariate with parameter $\boldsymbol{\beta}_x$ and with interaction effects with $t_{gk}$. From this model, genes with sum of squares of the test statistic for $\hat{\boldsymbol{\beta}}_x$ greater than a pre-defined threshold (see *chooseCandGenes(llStat.thrld)*) will be used as candidate genes for the next step. Among the genes that have a desired difference in the mean expression (obtained from Step 2), this step selects those with difference in the shape of the distribution.
- Step 4: from the set of genes identified in Step 3, it randomly selects $[p_{DE} \times G']$ genes($[.]$ = round to the nearest integer), which are designated as DE genes. For this set of genes, we estimate the density in (6) independently for each condition.
- Step 5: the rest of the $[(1 - p_{DE}) \times G']$ genes are designated as null genes, and we only use the data from one of the conditions for estimating the density and simulating new data for this set of null genes.

Steps 1 and 2 find genes with difference in mean expression more than the predefined threshold in the source data. Among these genes, steps 3 and 4 select those with difference in the shape of the distributions.

#### 1.5 Simulating batch effect

When samples/cells are generated from different instruments or cells come from different biological samples (for example subjects, tissue, etc), there will be a systematic variation which is called batch effect(Tung et al. 2017; Hicks et al. 2017). Given this information in the source data, our simulation procedure incorporates this batch effect in the simulated data if required. For designs with multiple batchs, we follow the following steps for each gene

- Step 1: for each batch $j = 1, \ldots, r$, we fit the model in (5)
- Step 2: for the estimated parameters of each batch obtained in step 1, we fit a multivariate normal distribution
- Step 3: to simulate a new data for a particular batch, we sample a parameter vector from the fitted multivariate normal distribution (in step 2), and use these parameters to construct the density in (6)

However, unless $r$ is sufficiently large ($r > 5$) for an accurate estimation of the multivariate normal distribution in step 2, we do not recommend this step.

### 2 Demonstration of SPsimSeq

#### 2.1 Datsets

For the subsequent demonstration and benchmarking of the SPsimSeq method, we use bulk and single cell RNA-seq datasets summarized below

- The Zhang data (L. Zhang and Zhang 2017): It was retrieved from GEO (accession number GSE49711) and contains 498 neuroblastoma tumors. In short, unstranded poly(A)+ RNA sequencing was performed on the HiSeq 2000 instrument (Illumina). Paired-end reads with a length of 100 nucleotides were obtained. To quantify the full transcriptome, raw fastq files were processed with Kallisto v0.42.4 (index build with GRCh38-Ensembl v85). The pseudo-alignment tool Kallisto (Bray et al. 2016) was chosen above other quantification methods as it is performing equally good but faster. For this study, we used the subset of the as used by Assefa et al. (2018) , i.e. a subset of 172 patients with high-risk disease were selected, forming two groups: the MYCN amplified ($n_1$ = 91) and MYCN non-amplified ($n_2$ = 81) tumors.

- Neuroblastoma NGP cells scRNA-seq data (NGP data): This data contains 83 NGP neuroblastoma cells (31 nutlin-3 treated and 52 controls). The data is generated using SMARTer/C1 protocol, and it was obtained from Verboom et al. (2018) study (GEO accession GSE119984).
- PBMC data: Contains 2700 single cells sequenced on an Illumina NextSeq 500 using unique moleculare identifiers (UMI). The data is generated using the 10x Genomics Chromium V2 protocol. The data is obtained from https://s3-us-west-2.amazonaws.com/10x.files/samples/cell/pbmc3k/pbmc3k_filtered_gene_bc_matrices.tar.gz

#### 2.2 Benchmarking: The Splat simulation method

To benchmark our simulation method, we implemented one of the most commonly used fully parametric simulation method, called the *Splat* simulation (Zappia, Phipson, and Oshlack 2017). This simulation uses a gamma-Poisson hierarchical model in which the mean expression level for each gene is sampled from a gamma distribution and the count for each cell is subsequently sampled from a Poisson distribution. Splat also adds outlier mean expressions and a mean-variance trend. Moreover, it uses a logistic function for the observed relationship between the mean expression of a gene and the proportion of zero counts to add excess zeros representing technical noise (aka dropouts). The Splat simulation is implemented using the *splatter* R Bioconductor package (version 1.6.1) (Zappia, Phipson, and Oshlack 2017). *Splat* uses simulation parameters learned from real data. To add a set of DE genes across simulated groups, it multiplies the mean expression of randomly selected genes by a factor, also known as a fold-change. To simulate bulk RNA-seq data using the Splat procedure, we disabled its feature that adds dropouts (*dropout.type="none"*), which is specifically designed for scRNA-seq data simulation (using the assay named *"true count"*, which does not add dropouts).

We compared the simulated data (using SPsimSeq and Splat) with the real source data with respect to various gene and cell/sample level metrics as used by Soneson and Robinson (2017) and Zappia, Phipson, and Oshlack (2017) to examine the quality of simulated data. These are,

- the distribution of mean and variance of the log-CPM across genes
- the relatioship between the mean and variance of the log-CPM of each gene

- the distribution of coefficients of variation (CV) in each gene and its relatioship with the mean log-CPM
- the distribution of the fraction of zero counts in each gene, and its relationship with the mean log-CPM.
- the distribution of the fraction of zero counts per cell/sample
- the distribution of library size across samples/cells
- the distribution of the correlation between samples/cells

#### 2.3 Simulation of bulk RNA-seq data

We use SPsimSeq to simulate bulk RNA-seq data starting from the Zhang neuroblastoma data. For benchmarking purpose we also simulate bulk RNA-seq data using the *Splat* procedures as described above. In particular, we simulate a bulk data with the following features

- 10,000 genes with 10% of them are differentially expressed
- a total of 180 samples
- samples are divided evenly into two groups (MYCN amplified and not amplified)
- all samples are generated in a single batch (as the Zhang data)

We use the following codes to generate the data according to the design listed above

```
# load required libraries
library(SPsimSeq)

# load the Zhang data (availabl with the package)
data("zhang.data")

# filter genes with sufficient expression (important step to avoid bugs)
zhang.counts <- zhang.data$counts[rowSums(zhang.data$counts > 0)>=10, ]
MYCN.status <- zhang.data$MYCN.status+1
```

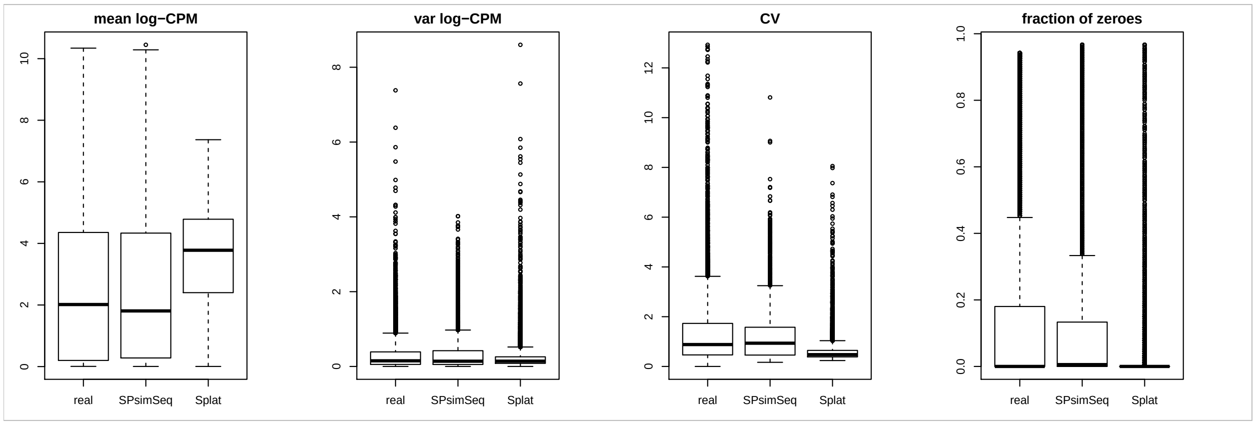

Figure 1: Comparing the distribution of mean log CPM, variance of log CPM, coeffcient of variation (CV), and fraction of zeroes per gene between the real Zhang data, SPsimSeq simulated data and the Splat simulated data.

```
# simulate data (we simulate here only a single data, n.sim = 1)
sim.data.bulk <- SPsimSeq(n.sim = 1, s.data = zhang.counts, batch = NULL,
                         group = MYCN.status, n.genes = 10000, batch.config = 1,
                         group.config = c(0.5, 0.5), tot.samples = 180, pDE = 0.1,
                         model.zero.prob = FALSE, result.format = "list", seed=25081988)
```

For the bulk RNA-seq simulation, the benchmarking results can be seen in Figure 1-3.

#### 2.4 Simulation of single cell RNA-seq data (read-count data)

Now we use SPsimSeq to simulate scRNA-seq data starting from the NGP neuroblastoma scRNA-seq data. We also simulate scRNA-seq data using the *Splat* procedures with dropouts added. In particular, we simulate a scRNA-seq data with the following features

- 10,000 genes with 10% of them are differentially expressed
- a total of 100 cells
- cells are divided evenly into two groups (nutlin-3 treated and control)
- all cells are processed in a single batch (as the data itself)

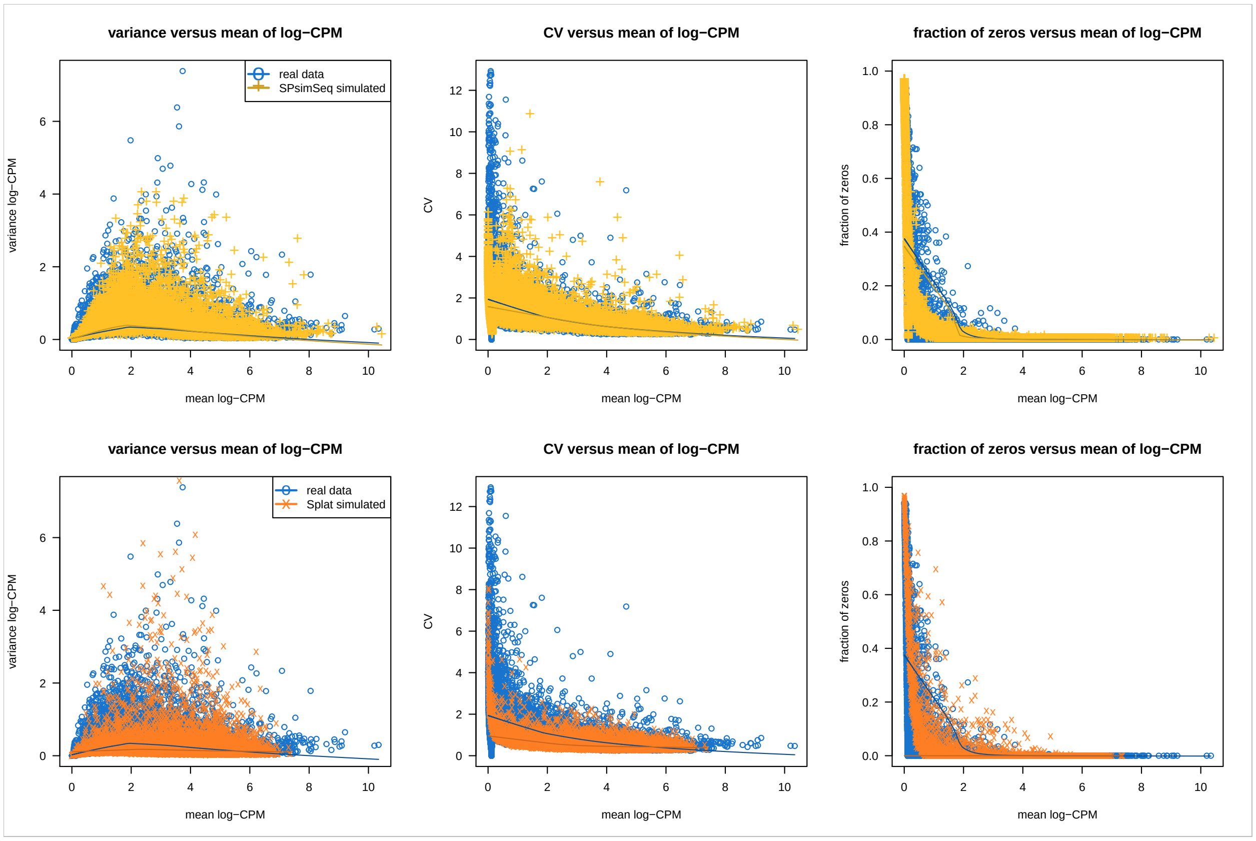

Figure 2: The relationship between mean and variance of log CPM, CV and mean of log CPM, and mean log CPM and fraction of zeroes per gene from the real data (Zhang data), SPsimSeq simulated data and Splat simulated data.

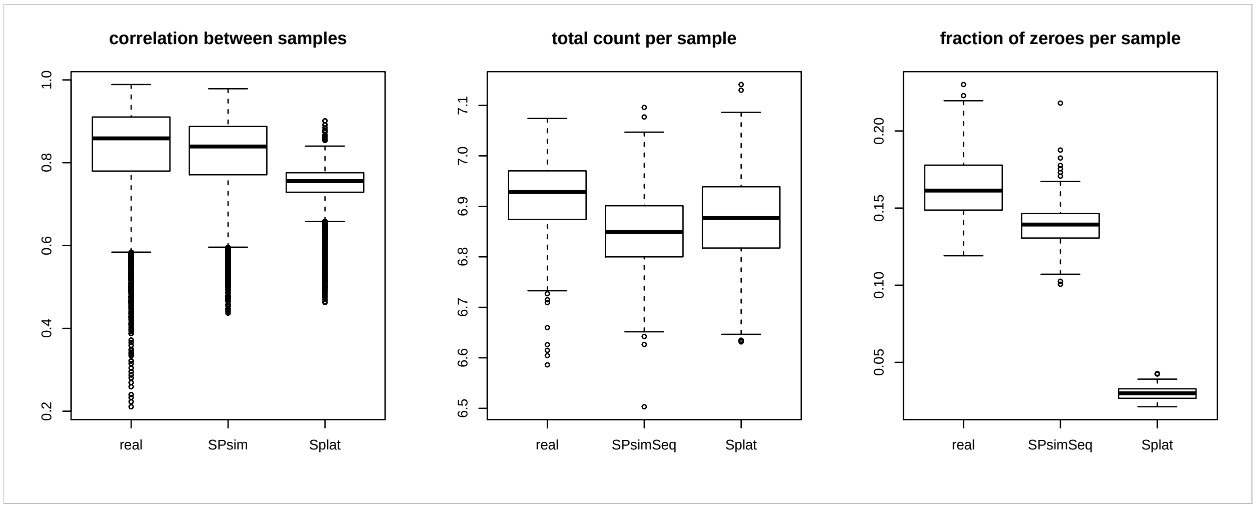

Figure 3: Distribution of the Pearson correlation coefficient between samples, total counts per sample (in log scale) and fraction of zeroes per sample from the real data (Zhang data), SPsimSeq simulated data and Splat simulated data.

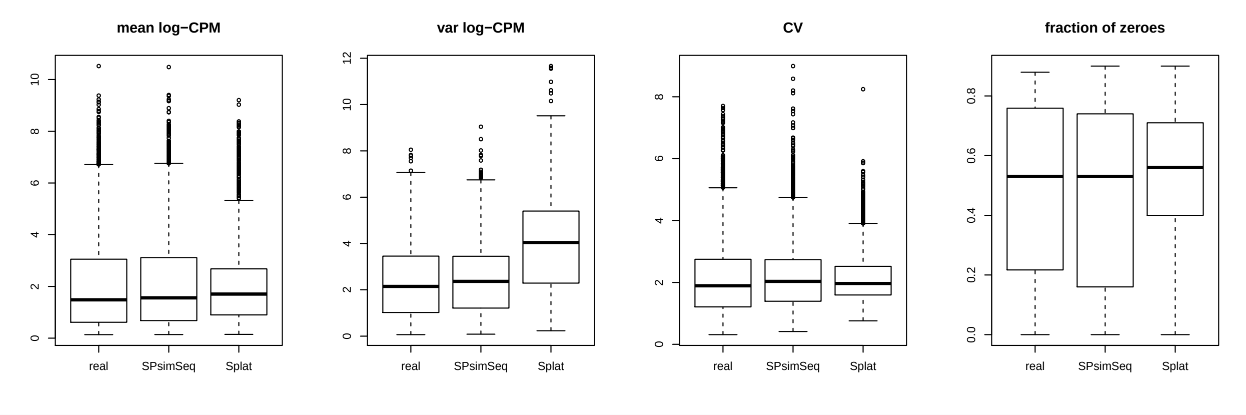

Figure 4: Comparing the distribution of mean log CPM, variance of log CPM, coeffcient of variation (CV), and fraction of zeroes per gene between the real scRNA-seq data, SPsimSeq simulated data and the Splat simulated data.

We use the following codes to generate the data according to the design listed above

```
# load required libraries
library(SingleCellExperiment)
library(SPsimSeq)

# load the NGP nutlin data (availabl with the package)
data("scNGP.data")

# filter genes with sufficient expression (important step to avoid bugs)
scNGP.data2 <- scNGP.data[rowSums(counts(scNGP.data) > 0)>=10, ]
treatment <- ifelse(scNGP.data2$characteristics..treatment=="nutlin",2,1)

# simulate data (we simulate here only a single data, n.sim = 1)
sim.data.sc <- SPsimSeq(n.sim = 1, s.data = scNGP.data2, batch = NULL,
                        group = treatment, n.genes = 10000, batch.config = 1,
                        group.config = c(0.5, 0.5), tot.samples = 100, pDE = 0.1,
                        model.zero.prob = TRUE, result.format = "SCE")
```

The benchmarking results for this simulation is presented in Figure 4-7.

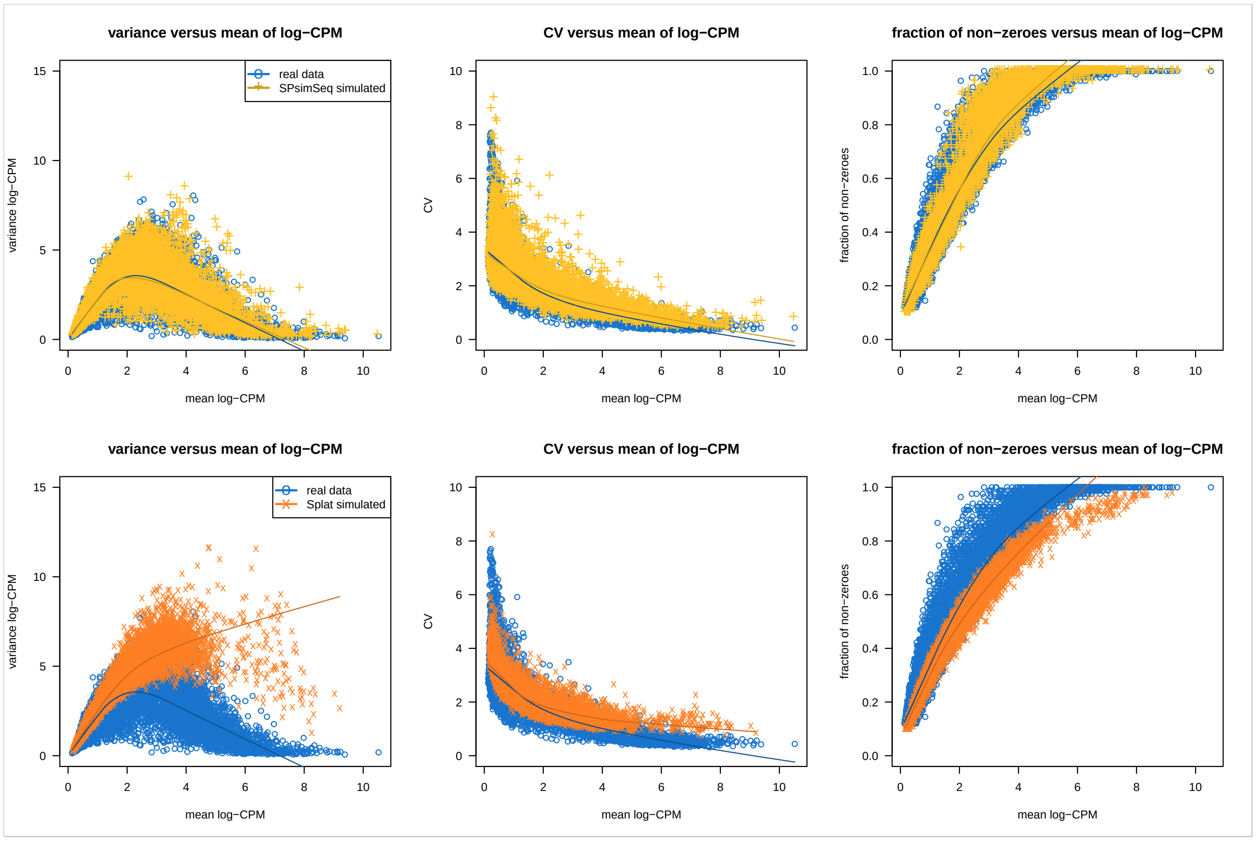

Figure 5: The relationship between mean and variance of log CPM, CV and mean of log CPM, and mean log CPM and fraction of zeroes per gene from the real C1 NGP scRNA-seq data, SPsimSeq simulated data and Splat simulated data.

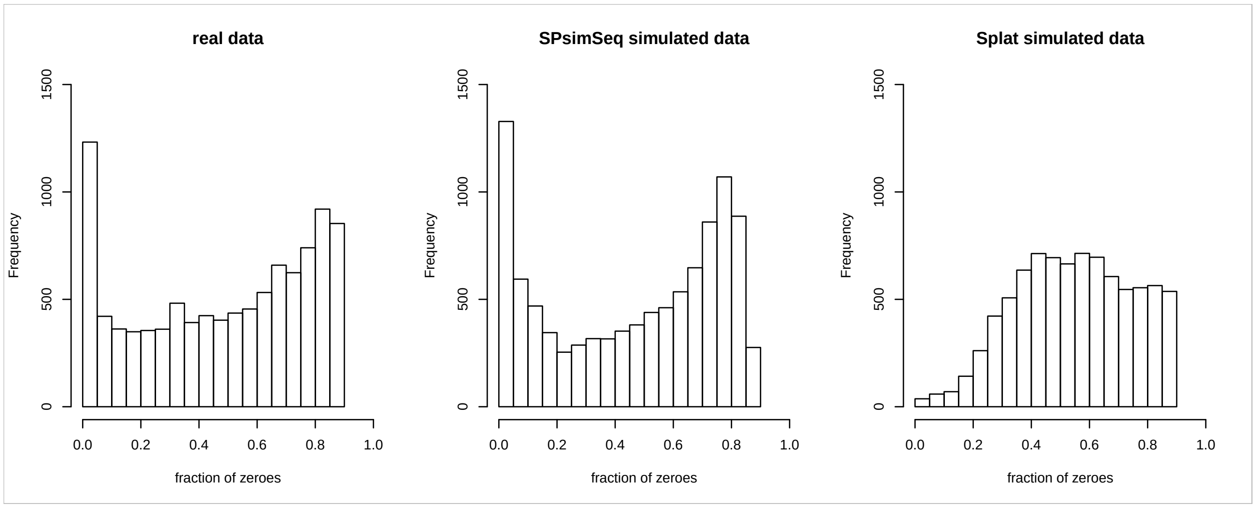

Figure 6: Distribution of the fraction of zero counts per gene from the real data, SPsimSeq simulated data and Splat simulated data. The boxplot version of this result is shown in Figure 4, the fourth panel

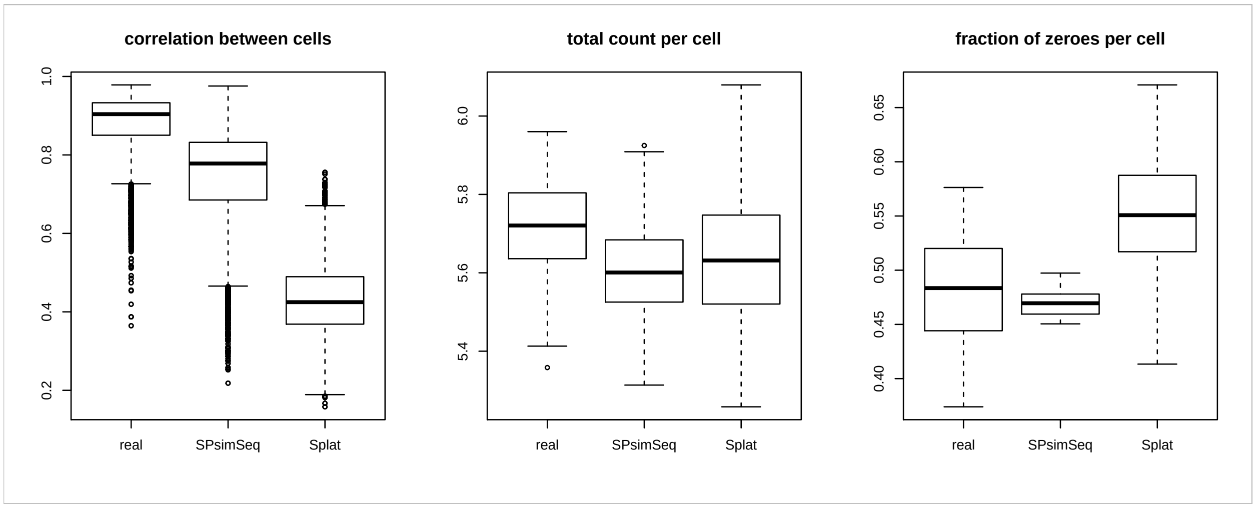

Figure 7: Distribution of the Pearson correlation coefficient between cells, total counts per cell (in log scale) and fraction of zero counts per cell from the real C1 NGP data scRNA-seq data, SPsimSeq simulated data and Splat simulated data.

#### 2.5 Simulation of single cell RNA-seq data (UMI-count data)

Unique molecular identifiers (UMI) are highly used in scRNA-seq studies. UMI counts reduce ammplification bias and result in better approximation of gene expression (Islam et al. 2014). In this section we demonstrate how SPsimSeq can be used to simulate UMI data for scRNA-seq studies. As a source data, we use the PBMC scRNA-seq data generated using Chromium protocol from the 10x Genomics, which contains UMI counts. We also use the *Splat* procedures to simulate UMI counts. In particular, we simulate a scRNA-seq data with the following features

- 10,000 genes
- a total of 500 cells
- all cells are from a single biological group
- all cells are processed in a single batch (as the data itself)

We use the following codes to generate the data according to the design listed above

```
# load required libraries
library(SingleCellExperiment)
library(SPsimSeq)
```

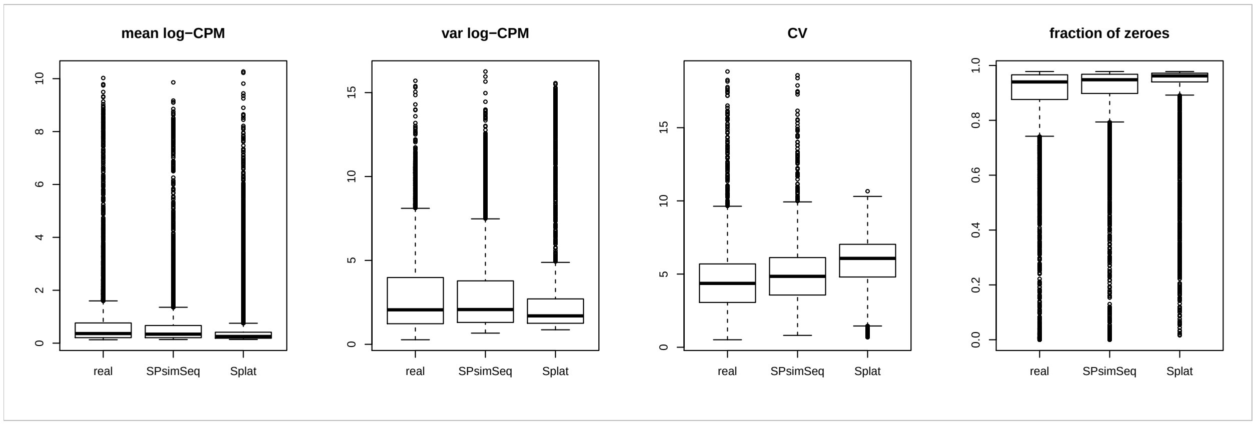

Figure 8: Comparing the distribution of mean log CPM, variance of log CPM, coeffcient of variation (CV), and fraction of zeroes per gene between the real UMI data, SPsimSeq simulated data and the Splat simulated data.

```
# load the Zhang data (availabl with the package)
data("PBMC.data")

# filter genes with sufficient expression (important step to avoid bugs)
PBMCdat2 <- PBMC.10x.data[rowSums(counts(PBMC.10x.data) > 0)>=20, ]

# simulate data (we simulate here only a single data, n.sim = 1)
sim.data.scUMI <- SPsimSeq(n.sim = 1, s.data = PBMCdat2, batch = NULL,
                           group = NULL, n.genes = 10000, batch.config = 1,
                           group.config = 1, tot.samples = 500, pDE = 0,
                           model.zero.prob = TRUE, result.format = "SCE")
```

The benchmarking results for this simulation is presented in Figure 8-11.

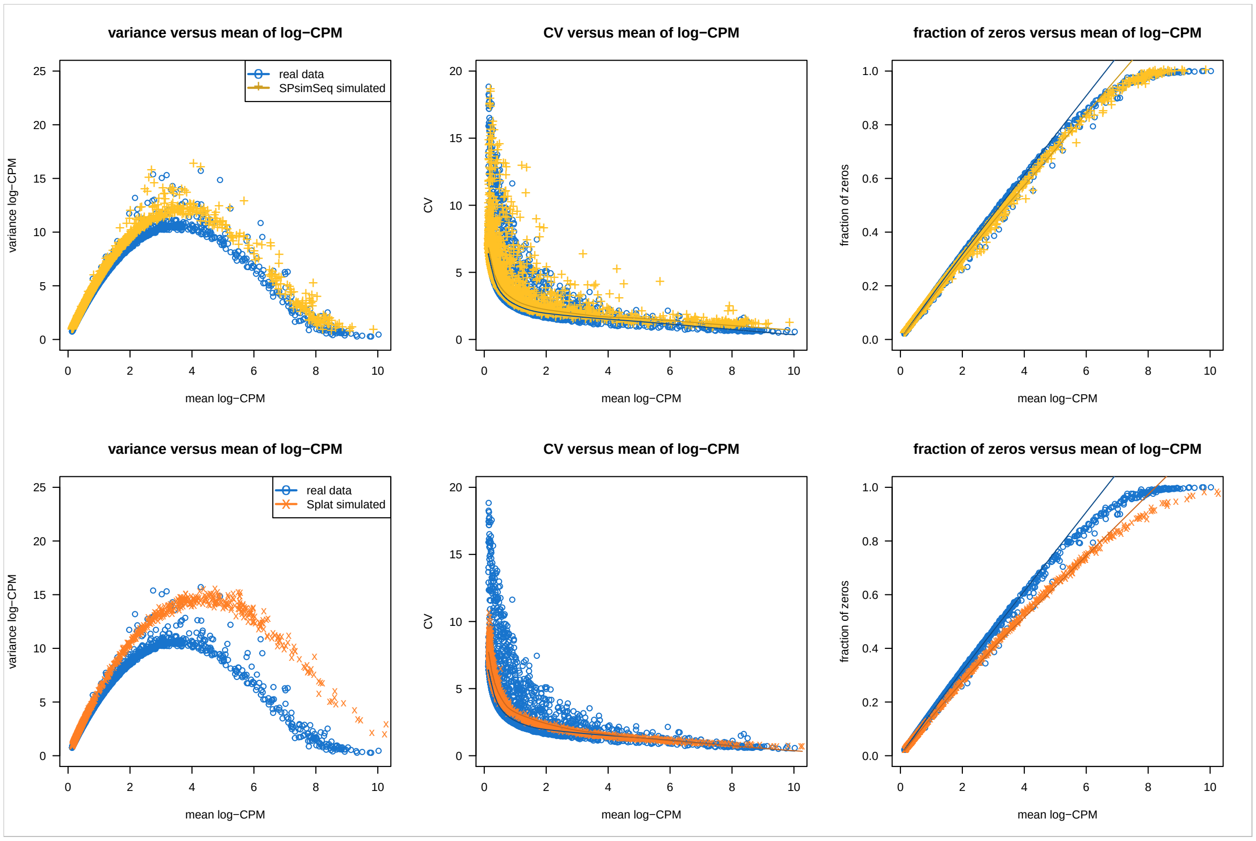

Figure 9: The relationship between mean and variance of log CPM, CV and mean of log CPM, and mean log CPM and fraction of zeroes per gene from the real UMI data, SPsimSeq simulated data and Splat simulated data.

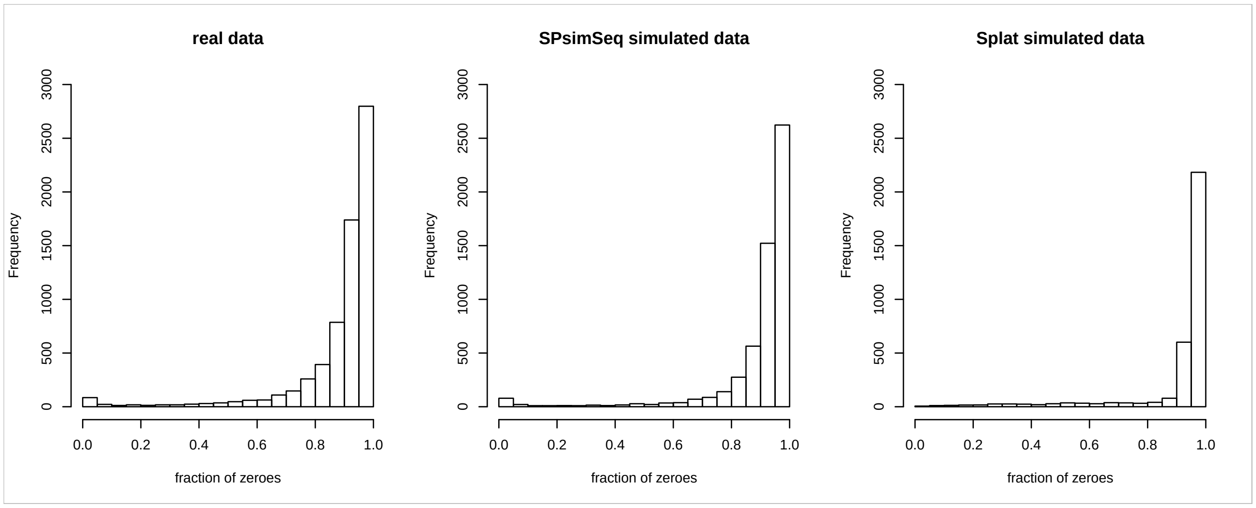

Figure 10: Distribution of the fraction of zero counts per gene from the real scRNA-seq data (UMI), SPsimSeq simulated data and Splat simulated data. The boxplot version of this result is shown in Figure 8, the fourth panel

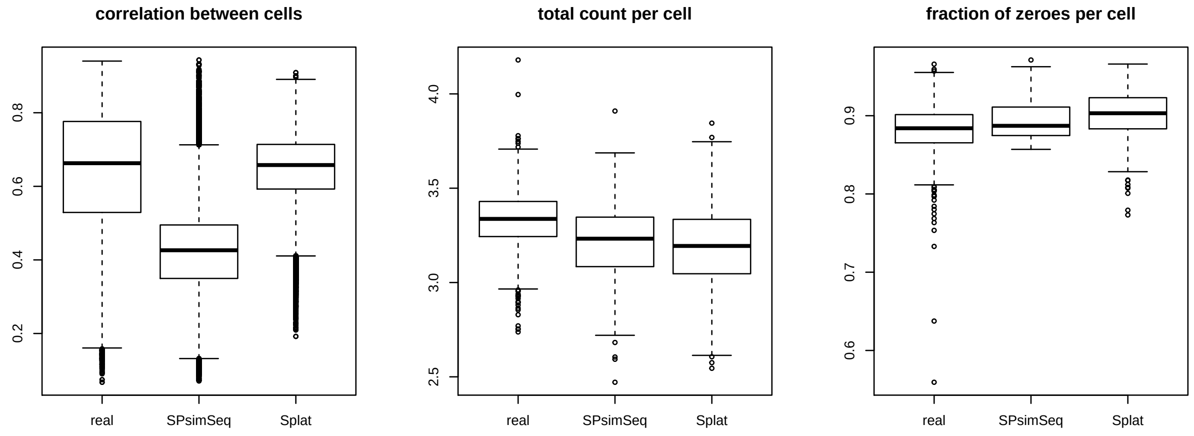

Figure 11: Distribution of the Pearson correlation coefficient between cells, total counts per cell (in log scale) and fraction of zero counts per cell from the real data scRNA-seq data (UMI), SPsimSeq simulated data and Splat simulated data.

Oxford University Press: 562–78.

Islam, Saiful, Amit Zeisel, Simon Joost, Gioele La Manno, Pawel Zajac, Maria Kasper, Peter Lönnerberg, and Sten Linnarsson. 2014. "Quantitative Single-Cell Rna-Seq with Unique Molecular Identifiers." *Nature Methods* 11 (2). Nature Publishing Group: 163.

R Core Team. 2018. *R: A Language and Environment for Statistical Computing*. Vienna, Austria: R Foundation for Statistical Computing. https://www.R-project.org/.

Soneson, Charlotte, and Mark D Robinson. 2017. "Towards Unified Quality Verification of Synthetic Count Data with countsimQC." *Bioinformatics* 34 (4). Oxford University Press: 691–92.

Sturges, Herbert A. 1926. "The Choice of a Class Interval." *Journal of the American Statistical Association* 21 (153). Taylor & Francis: 65–66. doi:10.1080/01621459.1926.10502161.

Tung, Po-Yuan, John D Blischak, Chiaowen Joyce Hsiao, David A Knowles, Jonathan E Burnett, Jonathan K Pritchard, and Yoav Gilad. 2017. "Batch Effects and the Effective Design of Single-Cell Gene Expression Studies." *Scientific Reports* 7. Nature Publishing Group: 39921.

Verboom, Karen, Celine Everaert, Nathalie Bolduc, Kenneth J Livak, Nurten Yigit, Dries Rombaut, Jasper Anckaert, et al. 2018. "SMARTer Single Cell Total Rna Sequencing." *bioRxiv*. Cold Spring Harbor Laboratory. doi:10.1101/430090.

Vieth, Beate, Christoph Ziegenhain, Swati Parekh, Wolfgang Enard, and Ines Hellmann. 2017. "PowsimR: Power Analysis for Bulk and Single Cell Rna-Seq Experiments." *Bioinformatics* 33 (21). Oxford University Press: 3486–8.

Zappia, Luke, Belinda Phipson, and Alicia Oshlack. 2017. "Splatter: Simulation of Single-Cell Rna Sequencing Data." *bioRxiv*. Cold Spring Harbor Labs Journals, 133173.

Zhang, Lihua, and Shihua Zhang. 2017. "Comparison of Computational Methods for Imputing Single-Cell Rna-Sequencing Data." *bioRxiv*. Cold Spring Harbor Laboratory. doi:10.1101/241190.
